# Automatic colocalization of high resolution MALDI MSI and Raman imaging applied to cardiac tissue of Fabry disease mouse models

**DOI:** 10.1101/2025.11.27.690907

**Authors:** Johann Dierks, Eike U. Brockmann, Anahi-Paula Arias-Loza, Thomas Bocklitz, Peter Nordbeck, Kristina Lorenz, Elena Tolstik, Sven Heiles

**Author notes:** Shared first authors (Johann Dierks, Eike U. Brockmann). Shared corresponding authors (Kristina Lorenz, Elena Tolstik, Sven Heiles).

## Abstract

Understanding early molecular changes in biological tissues is crucial for diagnosing pathological and genetic diseases and for elucidating their underlying mechanisms. However, localized molecular alterations of low molecular-weight compounds are not inferred from conventional staining or genetic methods. Here, we established a multimodal imaging approach that integrates Raman spectroscopy and atmospheric pressure matrix-assisted laser desorption/ionization mass spectrometry imaging (AP-MALDI MSI): two complementary, label-free techniques enabling molecular profiling of a broad spectrum of biomolecules from one single tissue section. This method was applied to detect Gb3 accumulation in heart tissue of murine models of Fabry disease, including mice deficient in α-galactosidase A (GLA) activity (GLA knock-out) and transgenic mice with a GLA knock-out and an upregulation of globotriaosylceramides (Gb3) synthase.

With AP-MALDI MSI we were able to discern the heterogenous expression of Gb3 lipoforms with down to 5 µm pixel size and reveal the significantly increased Gb3 content in mice containing a GLA knock-out combined with human Gb3 synthase overexpression compared to GLA knock-out and wild type samples. By employing Raman microscopy with a pixel size of 2 µm, we were able to contextualize the physiological alterations in cardiac tissue by identifying components associated with nuclei, tissue, collagen, and lipids for the same three genotypes. An automated co-localization algorithm aligned Raman and AP-MALDI-MSI data from the same tissue section with (5.1 ± 1.6) µm precision, enabling overlay at 5 µm and 2 µm resolution. The method resolved heterogeneous Gb3 distributions and distinct lipid species in cardiac mouse tissue.

## Introduction

Detection of molecular alterations in tissues can support early diagnosis, elucidate disease pathomechanisms, and facilitate timely implementation of targeted therapeutic options. However, the detection of such alterations in complex biological matrices – often with limited access and availability – remains a significant bioanalytical challenge, as comprehensive molecular profiling of lipid, metabolite, and protein distributions is essential for accurate disease predictions.^1–3^ Even though genetic tests are valuable diagnostic tools for mutation-driven diseases such as cancers or infiltrative cardiomyopathies, they have limited predictive power for disease onset or organ-specific manifestations as they do not capture the local microenvironments that critically influence these processes.^4,5^ Current diagnostic procedures analysing molecular changes rely on morphological, targeted immunohistochemical or biochemical assays lacking coverage of biomolecules across diverse chemical classes, especially those with low molecular weight.^6^

Fabry disease (FD) exemplifies these challenges as an X-linked lysosomal storage disorder caused by pathogenic variants in the gene encoding α-galactosidase A (GLA). These variants [can] result in reduced GLA activity and subsequent accumulation of globotriaosylceramide (Gb3) in diverse cell types.^7^ FD presents as a heterogeneous multisystemic condition ranging from multi-organ involvement to preferential cardiac or neurological involvement, with phenotypes varying significantly due to gender, genotype, age, and unidentified genetic modifiers.^8^ Despite the availability of genetic tests and assays for blood lyso-Gb3 and Gb3 levels,^9^ the individual and heterogenous deposition patterns of Gb3 make it difficult to infer the specific risk for kidney, brain, or heart damage and to determine optimal therapeutic strategies for individuals.^10^ To address these diagnostic limitations, spatially-resolved bioanalytic workflows offering untargeted molecular insights are needed as tools to provide additional information for risk stratification and therapeutic guidance.

Several micro-spectroscopy methods have been shown to be promising approaches for investigating untargeted local biomolecular changes in healthy versus diseased tissue such as Raman spectroscopy, coherent anti-Stokes Raman scattering (CARS) or stimulated Raman scattering (SRS) spectroscopy.^11,12^ By the detection of molecular fingerprints, these methods allow for the identification and spatial mapping of various molecular compound groups such as lipids, proteins, and nucleic acids as well as local accumulations of compounds like collagen with pixel sizes down to ∼ 400 nm.^13–16^ Thus, Raman imaging was employed to study lipid distributions in various cell types.^17,18^ Further, Hollon *et al.* used SRS to distinguish tumorous from healthy brain tissue in glioblastoma and guide intraoperative decision-making.^19^ Despite these potential applications in diagnosis and their analytical merits, the molecular identity of endogenous compounds, particularly low concentrated or heterogeneously distributed, is challenging, complicating the identification of biomarkers.^18^

Also, mass spectrometry imaging (MSI) methods such as matrix-assisted laser desorption/ionization (MALDI) provide detailed, spatially resolved information for many biomolecules in tissues. Atmospheric pressure (AP-) MALDI-MSI can detect changes in biomolecular profiles or distinct biomolecules, for example, distinguishing between healthy liver and granuloma.^20^ However, despite molecular specificity offered by MSI, the pixel resolution is often limited to 5 – 25 µm, and certain compound classes such as proteins are technically difficult to detect.^21,22^ Therefore, a bioanalytical workflow that combines molecular compound annotation via MSI with high lateral resolution and compound class specificity via Raman imaging is highly desirable to minimize analytical drawbacks and maximize information obtained from a single tissue section.

Indeed, previous proof-of-concept studies have demonstrated that Raman imaging and MSI provide complementary information when applied to biological tissues. The combination of Raman imaging and AP-MSI resolved histological features of mouse brain at lateral resolutions of 25 µm and 75 µm, respectively.^23–25^ Desorption electrospray ionization combined with Raman imaging enhanced molecular insights over Raman imaging or MSI alone in a multiple sclerosis mouse model.^26^ Streamlined pipelines such as RaMALDI have extended these approaches to diverse tissues including liver, kidney, and brain.^24^ Multimodal imaging clearly shows promise for capturing complementary molecular information, but key challenges remain. Achieving cellular-level resolution, and automated alignment of imaging methods is still difficult.

Here, we contribute to the field by combining high-resolution Raman imaging (2 µm spatial resolution) with AP-MALDI MSI (5 µm spatial resolution) in heart tissue from FD mouse models. Using automated image alignment, we achieved micrometre-level co-registration, enabling mapping of heterogeneous Gb3 accumulation together with nucleic acid, lipids, protein, and collagen distributions from one tissue slice. This approach not only enables higher spatial resolution for both modalities but also provides detailed molecular insights of FD-associated alterations in cardiac tissue, demonstrating the capabilities of hyperspectral multimodal imaging for the investigation of cardiac tissue.

## Materials and Methods

### Solvents and chemicals

Ammonium Acetate (≥97%, 238074, Sigma-Aldrich), Ammonium Formate (LiChropurTM, ≥99.0%, 70221, Supelco), Methyl-tert-butylether (MTBE, LiChrosolv®, 101845, Supelco), Acetone (LiChrosolv®, 100020, Supelco), Mayer’s Hematoxylin (MHS32-1L, Sigma-Aldrich), Eosin G (yellowish, 1.15935.0100), Ethanol (EMPLURA®, ≥99.5%, 818760, Supelco), 2-Propanol (used for matrix washing, ≥99.8%, 33539-M, Sigma-Aldrich), Eukitt® quick-hardening mounting medium (03989), SPLASH® Lipidomix® Mass Spectrometry Standard (330707, Avanti Polar Lipids LLC) and C17:0 Gb3 (860699P, Avanti Polar Lipids LLC) were all purchased from Merck KGaA, Darmstadt, Germany. Formic acid (LC-MS grade, 98%, 56302) was bought from Fluka, Honeywell International Inc, Charlotte, NC, USA. Methanol (Methanol absolute ULC/MS - CC/SFC, 136841), Water (Water ULC/MS – CC/SFC, 232141), 2-Propanol (2-Propanol ULC/MS – CC/SFC, 162641) and Acetonitrile (Acetonitrile ULC/MS – CC/SFC, 012041) were procured from Biosolve BV, Valkenswaard, NL. 2,5-Dihydroxyacetophenone (2,5-DHAP, ≥98%, A12185 was purchased from Alfa Aeser, Ward Hill, MA, USA. ROTI®Histol (6640.1) was bought from Carl Roth GmbH + Co. KG, Karlsruhe, Germany and Milli-Q water was produced by an ELGA PURELAB® flex 2 system (18.2 MΩ cm, ELGA LabWater Deutschland, Celle, Germany), 4% paraformaldehyd (PFA), Dulbecco’s phosphate buffered saline (PBS, Gibco, Thermo Scientific, Waltham, USA), commercial Gb3 mixture (Ceramide trihexoside, Cat. No. 1067, Matreya, USA).

### Mouse model

Male mice aged 18 to 20 weeks of a C57BL/6J genetic background were used for analysis. The genotypes are: GLA^WT^ mice, which represent the wild-type (WT) phenotype; hemizygous GLA^KO^ mice, which carry a targeted X-linked deletion of the α-Gal A gene resulting in a complete loss of enzyme activity^27^ and G3Stg/GLA^KO^ mice, a model that combines the α-Gal A knockout with the expression of human Gb3 synthase.^28^ All animal procedures were conducted in compliance with German animal welfare regulations and approved by the regional authority for animal research (Landesamt für Verbraucherschutz und Ernährung NRW, Germany; approval number Az. 81-02.04.2021.A464).

### Cryosectioning

Heart tissues were sectioned into 20 µm thick sections at −22°C using a cryotome (Thermo Scientific, Waltham, USA) after attachment of tissues with ice water to the cryotome holder. Resulting sections were placed on CaF_2_ slides of 2 mm thickness (Korth, Kiel, Germany) or microscopic glass slides (Th. Geyer GmbH & Co. KG, Renningen, Germany). The slides were stored at −80 °C until further use.

### Raman data acquisition

On the day before the Raman measurements, samples were moved from −80 °C to −20 °C for slow thawing. The sections were fixed at RT in 4% PFA solution for 10 min, washed with PBS and afterwards vacuum-dried (Concentrator plus, Eppendorf SE, Hamburg, Germany) for 15 min. Several red marks were added around the tissue sections for Raman/MALDI colocalization.

Raman images were acquired using a confocal Raman microscope (alpha 300R, WITec, Ulm, Germany). Brightfield (BF) images were acquired for both tissue overview and selected regions of interest. For all Raman measurements, a 785 nm excitation laser beam was focused on the sample using a 50x/0.75 NA dry objective (Zeiss, Jena, Germany). A laser power of 100 mW was used, and the integration time was set to 2 s. The pixel resolution of all images was 2 µm. A 300 g/mm grating was used for data acquisition.

For reference spectra, a commercial Gb3 mixture was dried and measured in single point measurements using a 785 nm laser source with 50 mW power, 30 s integration time and 10 accumulations.

### Raman data analysis

Data analysis of all measurements was performed using in-house written algorithms implemented in python 3.12 using numpy 1.26^29^, scipy^30^ and scikit-learn^31^ for computation and matplotlib^32^ for visualization. First, the data were pre-processed using an automatic cosmic spike correction^33^, wavenumber calibration to a 1 cm^-1^ grid, baseline correction using sensitive iterative peak (SNIP) clipping of the pybaseline package^34,35^, Savitzky-Golay smoothing, background segmentation using k-means clustering, and vector normalization.

After pre-processing, images were analysed individually using a combination of hierarchical cluster analysis (HCA) and a multi-curve resolution with alternating least squares (MCR-ALS). First, the measurements were clustered individually with Ward algorithm and Euclidean distance metric into 10 or 100 clusters, respectively. Afterwards the mean spectra of all images were merged into one single dataset and MCR-ALS was performed to detect important chemical components. MCR-ALS aims to reconstruct this data analysis matrix D as follows:

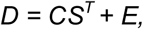

where C denotes the concentration or abundance matrix, ST the transposed spectral component matrix and E the residual errors, which are used to describe each pixel spectrum and to create abundance distribution.^36^ MCR-ALS was applied with the pymcr 0.5.1 package^37^ with Ordinary Least Square regressors for both the concentration and the components. Constraints were set to nonnegativity for components and nonnegativity plus normalization to 1 for concentrations. The number of components was manually varied and the minimal number with an appearance of a lipid-associated component was used for final analysis. Spectra were initialized using classical k-means clustering with default settings of the scikit-learn package. For statistical analysis, components were tested individually computing the mean abundance per mouse based on the tissue sections. Normal distribution was tested with Shapiro-Wilk test and Levene-test was used for homogeneity testing. If successful, one-way ANOVA with Bonferroni correction was applied and – if successful – a post-hoc Tukey test to detect group-specific significances.

### Sample preparation for AP-MALDI MSI

For AP-MALDI MSI sample preparation, sections on glass slides were thawed and dried in a desiccator for 30 min. Matrix was applied to the samples using a custom-built sublimation chamber (ChemGlass). A total of 250 µL of a 20 mg/mL 2,5-DHAP solution in acetone was added to the sublimation chamber and the acetone was completely evaporated. Vapour deposition was performed at 130 °C for 4 min under vacuum. The sample was cooled with water ice to 0.5 °C. To prevent recrystallization due to humidity, the sample was immediately transferred to a desiccator for 30 min. As quality control measure, BF images of the sample were recorded before matrix application, after matrix application, and after AP-MALDI MSI acquisition.

### AP-MALDI MSI data acquisition

AP-MALDI MSI measurements were conducted on a QExactive™ HF orbital trapping mass spectrometer (Thermo Fisher Scientific, Bremen, Germany) equipped with an AP-MALDI^5^ AF ion source (TransMIT, Gießen, Germany). Samples were analysed at a pixel resolution of 25 x 25 µm^2^ (full pixel mode) and 5 x 5 µm^2^ (spot mode).^38^ The global 20% attenuator was activated, and the variable attenuator was set to 38°C. Data were acquired in positive-ion mode with a scan range of *m*/*z* 200-2000 for 25 µm measurements and *m*/*z* 500-1500 for 5 µm measurements. The mass resolution was set to 240,000, automatic gain control was disabled, the maximum injection time was 500 ms. The acceleration voltage was set to 3.5 kV for 25 µm measurements and 2.5 kV for 5 µm measurements. The capillary temperature was maintained at 275 °C, and the S-lens RF level was set to 100.

### AP-MALDI MSI Data analysis

Thermo .RAW-files and TransMIT .udp-files were converted to .imzML format using the RAW+UDP to IMZML Converter (v1.8r3, TransMIT, Gießen, Germany). The resulting .imzML files were processed using LipostarMSI (v2.1.0b7, Molecular Horizon, Bettona, Italy).^39^ Data were recalibrated based on the lock masses *m*/*z* 826.5723 (PC(36:1), [M+K]^+^) for 25 µm measurements and *m*/*z* 756.5514 (PC(32:0), [M+Na]^+^) for 5 µm measurements. The recalibrated data were exported as .imzML-files and uploaded to Metaspace^40^ (https://metaspace2020.org) for preliminary lipid annotation.

For statistical analysis of Gb3 accumulation in cardiac tissue, five AP-MALDI MSI datasets for each genotype were loaded into a single LipostarMSI session and normalized to the root-mean-square (RMS) intensity. Normalized data were segmented using bisecting k-means clustering and spatial-aware processing. ROIs covering the tissue section were defined based on the previous segmentation. The mean intensities for the molecules of interest were then extracted and processed in Prism (v10.2.3, GraphPad Software, LLC, Boston, MA, USA). For statistical analysis, normal distribution was tested with the Shapiro-Wilk test and the Grubbs method was used for the identification of potential outliers. Variance analysis was performed using a one-way ANOVA with Bonferroni correction. Since all Gb3 species in the GLA^WT^ group were below the detection limit, resulting in mean intensities of zero, this group was excluded from statistical analysis due to the lack of normal distribution and variance. Consequently, an unpaired t-test was conducted for the comparison between the remaining two groups. Normal distribution was tested with the Shapiro-Wilk test and the Grubbs method was used for the identification of potential outliers. Variance analysis was performed using a one-way ANOVA with Bonferroni correction. For this we assume that the variance between groups is bigger than the biological variance within a group. Since all Gb3 species in the GLAWT group were below the detection limit, resulting in mean intensities of zero, this group was excluded from statistical analysis due to the lack of normal distribution and variance. Consequently, an unpaired t-test was conducted for the comparison between the groups in which the mean of the lipid intensity was not zero.

The nomenclature for the lipid species follows the LIPID MAPS shorthand notation guidelines (https://www.lipidmaps.org/lipid_nomenclature).^41,42^

### Overlay of AP-MALDI MSI and Raman measurements

For the combination of both measurement modalities, an inhouse-written application in python 3.12 utilizing the opencv (https://opencv.org/) and scikit-image 0.25.2 packages was developed.^43^ First, Raman measurements were inserted at their positions in the Raman BF overview, which included the whole tissue section with red markers based on coordinates retrieved from system metadata. The metadata were automatically delivered by the system after each Raman measurement and included the x, y position in the coordinate system. Afterwards, a pattern matching algorithm between BF overview image and BF taken in advance of each Raman measurement was applied to optimize the position. Pattern matching was achieved using a fast normalized cross-correlation algorithm (implemented via the match_template function of the scikit-image package) with variation of x, y position. The region of interest to detect the optimal position was set to 1 mm^2^ region around the position determined by the metadata. The position of maximum response was computed and stored as final coordinate.

In a second step, the AP-MALDI MSI measurements were localized in the MALDI-associated BF overview (that was recorded after the measurements). Edge detection maps were computed for each BF overview image using a holistically nested edge-detection neural network.^44^ The network was not pretrained on the images and integrated using the dnn function of opencv package. Images were forward passed through the network and edge detection map was used for ongoing computations. Detection of rectangular structures was achieved by mask convolution on the BF overview image. The mask was designed based on pixel size of the AP-MALDI MS measurement and ratio between AP-MALDI MSI resolution and BF overview resolution, where the estimated measurement border was set to 255 with a thickness of 5 pixels. The outer and inner edges were set to 1. The remaining pixels were set to 0 excluding them from computation. Signal of maximum response was computed to determine the optimal position. This process was repeated twice, first just by varying the mask position in the image to determine a general region of interest and afterwards varying the mask position and angle in this ROI.

The final step was the coalignment of the BF overview images and consequently all measurements. Red markers were detected in both images by converting the image to HSV colour space, just keeping the saturation dimension (which is far higher for artificial marker pens compared to biological tissue) and applying Otus’s thresholding to extract them. Afterwards, morphological operations in form of hole closing and removing small objects below 10 µm were applied. Then, the affine matrix describing translation, rotation and scaling was computed between the BF overviews by applying the cv2.estimateAffinePartial function. For Raman imaging and MSI overlays, 5 µm AP-MALDI MSI pixels were scaled to 2 µm pixels using Nearest-Neighbour interpolation to display these two modalities in one singular image. This matrix was stored and used for all coordinate translation.

After co-registration, overlapping regions were determined by coordinate comparison. For merging, the MCR-ALS results of the Raman analysis were overlapped with the AP-MALDI MSI specific results of Gb3. To determine if each pixel signal per component is signal or background, classical Otsu’s thresholding was applied for each component, and the signal was merged into one single image.

## Results and Discussion

### Study Design

In order to investigate molecular changes during disease development, we used three FD mouse models, including wild-type, GLA^KO^ and G3Stg/GLA^KO^ mice. The wild-type (GLA^WT^) mice served as control group, GLA^KO^ mice, lacking GLA, as model for early stage or asymptomatic FD phenotype with moderate Gb3 accumulations^27^, and G3Stg/GLA^KO^ mice, GLA^KO^ mice combined with overexpression of human Gb3 synthase in order to further increase Gb3 depositions^28^, as a model for a symptomatic FD phenotype including renal and cardiac dysfunction.^28,45^

First, aiming to establish a comprehensive workflow for the implementation of AP-MALDI MSI and Raman imaging and their simultaneous application, we started with separate measurements on consecutive sections of cardiac tissue (left ventricular mouse heart sections) by Raman imaging and AP-MALDI MSI. These parallel measurements provided information for localization of biochemical components in the cardiac tissue along with specific molecular identification and spatial distribution analysis of distinct Gb3 species. Next, we performed sequential Raman imaging and AP-MALDI MSI imaging using the same tissue sections and established comprehensive multimodal overlays of molecular images from both modalities. This overall procedure aims to maximize the combined molecular information, to differentiate between the three genotypes and to achieve an automated overlay of Raman imaging and AP-MALDI MSI modalities.

### AP-MALDI MSI of Gb3 in murine cardiac tissue

First, we aimed to determine the detection capability of AP-MALDI MSI for Gb3 accumulation in heart tissue across the three experimental groups: GLA^WT^, GLA^KO^, and G3Stg/GLA^KO^ mice. For this purpose, heart tissue sections were scanned with 25 µm step size and statistical analysis was employed for ROIs covering the entire tissue section of each AP-MALDI MSI measurement. **Figure 2A** shows BF images of heart tissue sections alongside corresponding AP-MALDI MSI measurements and post-acquisition H&E staining, displaying spatial intensity distributions for Gb3(34:1) (*m*/*z* 1062.635; [M+K]^+^), Gb3(42:2) (*m*/*z* 1172.744; [M+K]^+^), and PC(34:1) (*m*/*z* 798.542; [M+K]^+^). **Figure 2B** contains exemplary single pixel mass spectra for each group with the respective Gb3 lipoforms annotated based on the total number of carbon atoms and C=C double bonds in the FA chains. The statistical analysis of ROIs is shown in **Figure 2C** as violin plots for Gb3(34:1), Gb3(42:2) and PC(34:1).

**Figure 1:**
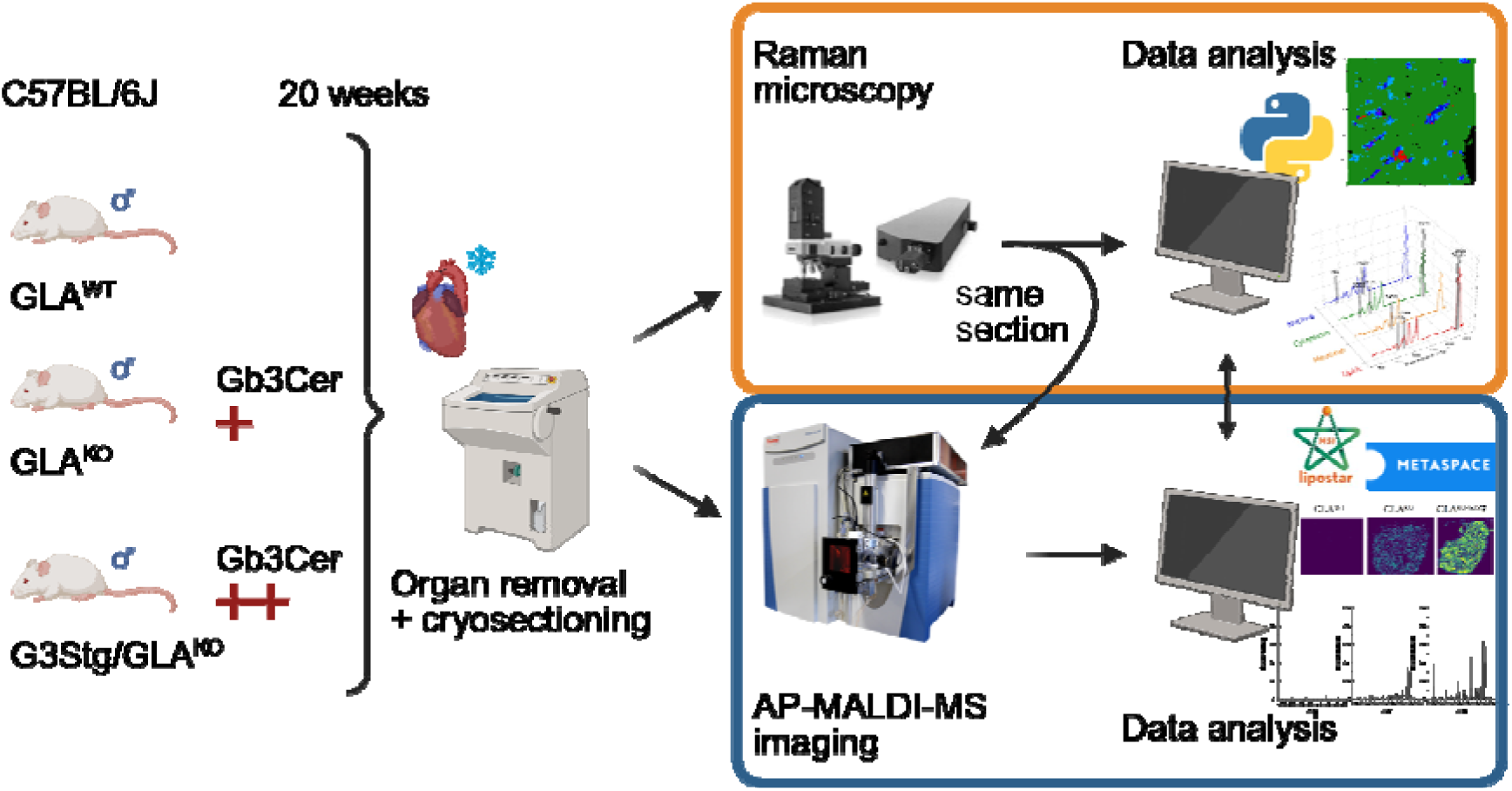
Study concept. Determination of cardiac Gb3 accumulation in a FD mouse model, consisting of wild-type (GLA^WT^), GLA knockout (GLA^KO^), and knockout with additional human Gb3 synthase expression (G3Stg/GLA^KO^) mice, was performed on heart cryosections from 20-week-old animals. Initially, AP-MALDI MSI and Raman microscopy measurements were conducted on consecutive sections. Subsequently, both techniques were applied to the same section, first using Raman imaging, followed by AP-MALDI MSI, and data analysis. The resulting datasets from both modalities, namely data of consecutive sections and of sections with combined measurements, were integrated to enhance the depth of information. Created in https://BioRender.com.

**Figure 2:**
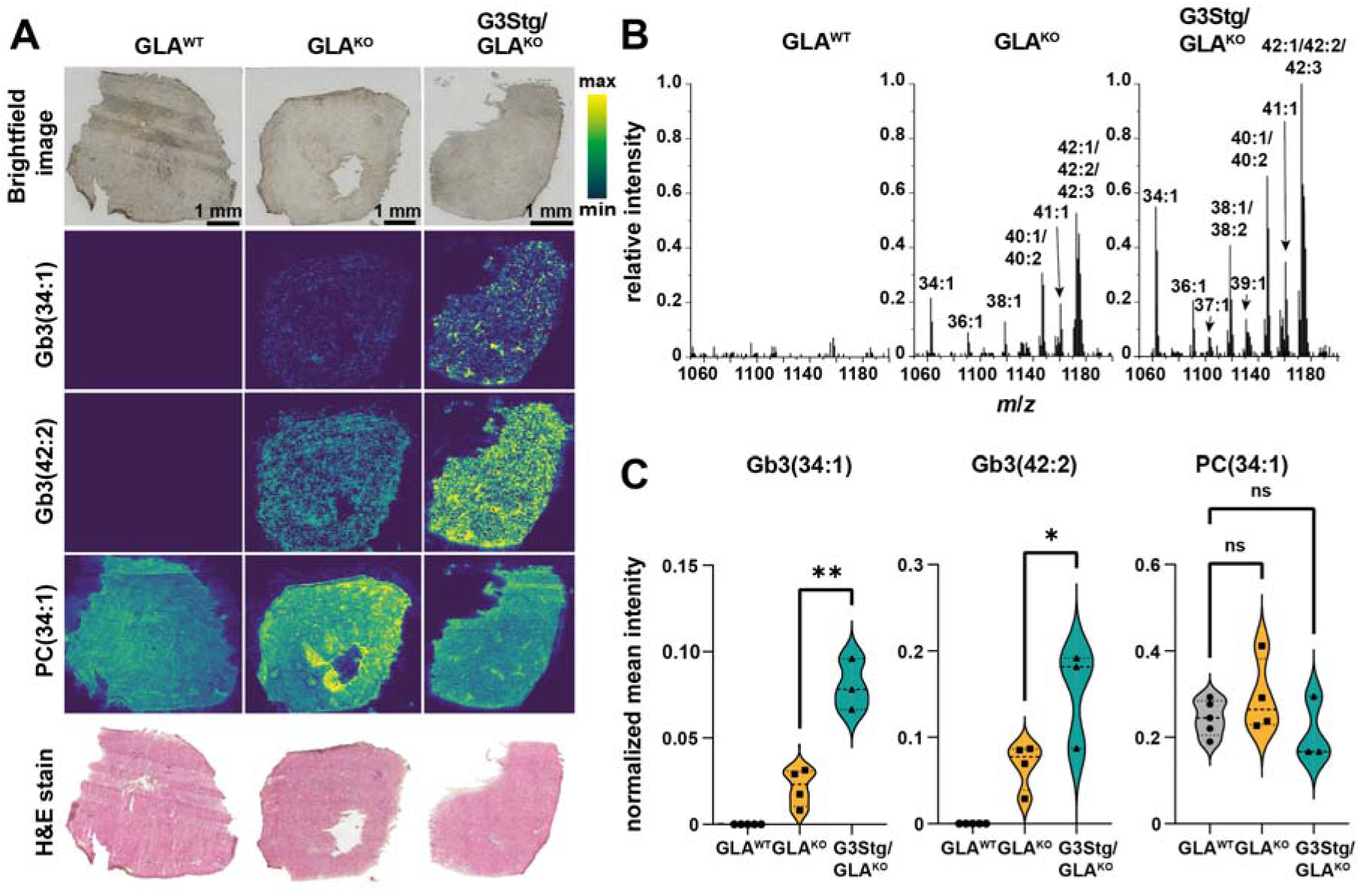
Gb3 accumulation from AP-MALDI MSI measurements. **A)** Visualization of Gb3 accumulation in murine cardiac tissue of the GLA^KO^, G3Stg/GLA^KO^ and GLA^WT^. Exemplary intensity distributions are presented for Gb3(34:1) (m/z 1062.6345; [M+K]^+^), Gb3(42:2) (m/z 1172.74380; [M+K]^+^) and PC(34:1) (m/z 798.5418; [M+K]^+^). Data were acquired with a pixel size of 25 x 25 µm^2^ in full pixel mode and RMS-normalized. **B)** Comparison of representative single-pixel mass spectra in the range from m/z 1050 to 1200. Annotation was performed based on the accurate mass of [M+K]^+^-adducts and MS/MS analysis. **C)** Statistical analysis of the Gb3 accumulation was conducted using AP-MALDI MSI measurements of murine cardiac tissue. The displayed intensities represent mean ROI intensities of the tissue area and were calculated following hot-spot removal by histogram equivalation and RMS normalization. GLA^WT^ n=5, GLA^KO^ n=4, G3Stg/GLA^KO^ n=3 biological replicates. **, p < 0.01; *; p < 0.05; ns = not significant.

Based on the colour-coded intensity maps in **Figure 2A**, a pronounced Gb3 accumulation in tissue of G3Stg/GLA^KO^ mouse hearts relative to GLA^KO^ samples was measured. Notably, Gb3 signals were absent in GLA^WT^ control tissue. In addition to these genotype-dependent differences in Gb3 accumulation, we observed heterogeneity in the fatty acyl chain composition of accumulated Gb3 species. This is exemplified by Gb3(34:1) and Gb3(42:2), which contain a C16:0 and a C24:1 fatty acyl chain, respectively. However, their distribution within the heart tissue was not identical, varying in intensity and localization (**Figure 2A**). Additional Gb3 lipoforms identified based on AP-MALDI MSI and LC-MS/MS analysis are summarized in **Figure S1** and **S2,** and **Table S1**. The ubiquitous lipid PC(34:1) served as an internal quality control exhibiting no statistically significant alterations between genotypes and the identification of tissue outlines as well as the conformation of comparability of MSI datasets.

Using AP-MALDI MSI, we detected and annotated 12 distinct Gb3 species in G3Stg/GLA^KO^ samples and nine in GLA^KO^ samples, whereas no Gb3 species were detected in GLA^WT^ samples (**Figure 2B**). Validation of the annotated lipids detected by AP-MALDI MSI was performed using LC-MS/MS of pooled lipid extracts, which identified 20 Gb3 species based on characteristic MS/MS fragmentation patterns (**Table S1**). This dual approach confirmed the molecular identity of the AP-MALDI MSI annotations but also its capability to capture the spatial resolution of predominant Gb3 variants in the tissue.

Not only Gb3(34:1) and Gb3(42:2) revealed a significantly enhanced accumulation of these disease specific lipid species in myocardial Fabry samples of the G3Stg/GLA^KO^ genotype compared to GLA^KO^ (**Figure 2C**) but also other Gb3 compounds were significantly upregulated (**Figure S1**).

Our data confirm the accumulation of Gb3 in the FD mouse model with elevated Gb3 intensities in the GLA^KO^ samples due to the hindered degradation of Gb3 and even higher intensities in the G3Stg/GLA^KO^ samples which can be explained by a higher production of Gb3 through the insertion of G3Stg. Through our refined AP-MALDI MSI method we visualized the distribution of the most abundant Gb3 lipoforms as described in the literature.^46^ In the GLA^WT^, Gb3 signals were absent or below the limit of detection (LOD) of our AP-MALDI MSI method. Of note, we could not detect lyso-Gb3 and analogues in heart tissue of FD mice (GLA^KO^ or G3Stg/GLA^KO^) using AP-MALDI MSI, consistent with lyso-Gb3 concentrations having been reported about 50 times lower than Gb3 concentrations in human heart tissue homogenates.^47,48^

To assess whether the heterogeneous distribution of Gb3 in myocardial tissue is dependent on the Gb3 lipoform and if it correlates with distinct cell types, we conducted high resolution AP-MALDI MSI measurements with higher spatial resolution and streamlined the workflow for multimodal imaging.

### Determination of heterogeneous Gb3 lipoform accumulation

**Figure 3A** shows heterogenous Gb3 distribution across murine cardiac tissue, with accumulation appearing as localized intensity spots. To improve the resolution of our AP-MALDI MSI results, we conducted AP-MALDI MSI at 5 x 5 µm^2^ pixel resolution on PFA-fixed tissue. The intensity maps acquired by high resolution AP-MALDI MSI highlights the magnitude and heterogenous spatial localization between Gb3 molecular lipoforms (**Figure 3A**). Whereas some of the Gb3 signals exhibit pronounced accumulation in selected regions of the tissue as highlighted by arrows in **Figure 3A**, Gb3 compounds differing solely in the fatty acyl composition have lower signal intensity in these respective regions and increased signal intensities in other areas of the tissue. This is best represented by the overlay image for selected examples shown in **Figure 3B**. Here, intensity maps of the differential spatial patterns of Gb3(42:2) ([M+Na]^+^, *m*/*z* 1156.769, orange), Gb3(38:1) ([M+Na]^+^, *m*/*z* 1102.722, blue) and the omnipresent lipid PC(34:1) ([M+Na]^+^, *m*/*z* 782.567, grey) are overlaid. Comparison to the BF and H&E images suggests that these regions of Gb3 accumulations are confined to one single or few neighbouring cells in cardiac tissue (**Figure 3B**).

**Figure 3:**
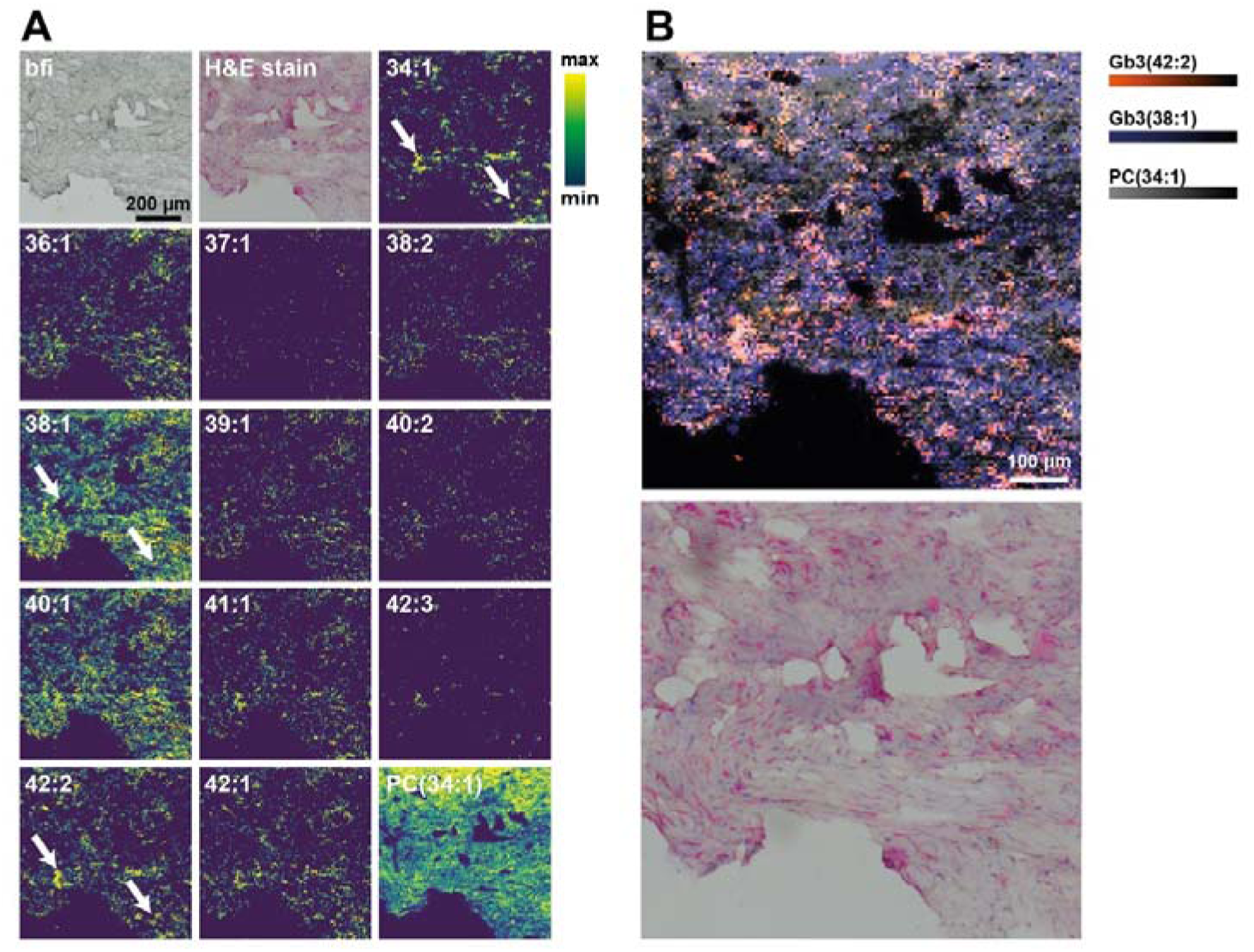
High-resolution AP-MALDI MSI intensity maps of Gb3 species in a G3Stg/GLA^KO^ sample. **A)** Shown are BF images of unstained and H&E stained tissue and the intensity distributions of all detected Gb3 species as [M+Na]^+^-adducts. Data were recorded after formalin fixation and Raman microscopy acquisition from CaF_2_-glass discs with 5 x 5 µm^2^ pixel size in spot acquisition mode. Annotation was performed based on the accurate mass of [M+Na]^+^-adducts and MS/MS analysis. **B)** Overlay of Gb3(42:2) ([M+Na]^+^, *m*/*z* 1156.7691, orange), Gb3(38:1) ([M+Na]^+^, *m*/*z* 1102.7223, blue) and PC(34:1) ([M+Na]^+^, *m*/*z* 782.5673, grey) and the corresponding bright-field image of H&E stained tissue.

Our high resolution MALDI-MSI data, visualizing the different intensity distributions based on Gb3 lipoforms, are in line with previous findings by Onoue *et al.* where they manually correlated the localization of Gb3 species with vacuolar degeneration in cardiomyocytes of human endocardial biopsy samples.^49^ The heterogeneous accumulation of Gb3 in the heart tissue of FD models is known from Gb3 staining and has been described for different cardiac cell types such as cardiomyocytes, endothelial cells and smooth muscle cells of vessels, conduction tissue or fibroblasts.^10,49–54^ However, the distinct molecular composition and the heterogeneity between these lipid compounds is not available from staining experiments. Despite the existence of several studies that employed MALDI MSI for the characterization of Gb3 accumulation in FD in a variety of tissues including kidney, heart, and skin^49,55–58^, until now no data with comparable degree of detail and spatial resolution for the determination of FD associated Gb3 accumulation in heart tissue is available.

### Raman imaging of heart tissue from murine FD model

For Raman microscopy, the images of mouse heart sections of the three genotypes were acquired and analysed first applying HCA analyses (**Figure 4**). The number of HCA clusters set to 10 allowed a separation of DNA/nucleic acids accumulation, typically associated with nuclei or mitochondria, and the tissue matrix (**Figure 4A**). The “tissue” clusters were characterized by Raman intensity peaks at 1003 cm^-1^ representing phenylalanine, at 1255 cm^-1^ representing amid III band, and at 1445 cm^-1^ representing CH_2_/CH_3_ stretching.^59,60^ The DNA/nucleic acids (“nucleus”) clusters consisted of Raman intensity peaks at 785 cm^-1^ and 1095 cm^-1^ characteristic for DNA backbone and nucleic acids.^59^ The localization of these clusters is presented on the colour maps (**Figure 4C, left column**; “nucleus” in blue; “tissue” in green). The distribution and number of pixels of these clusters were comparable in all genotypes.

**Figure 4:**
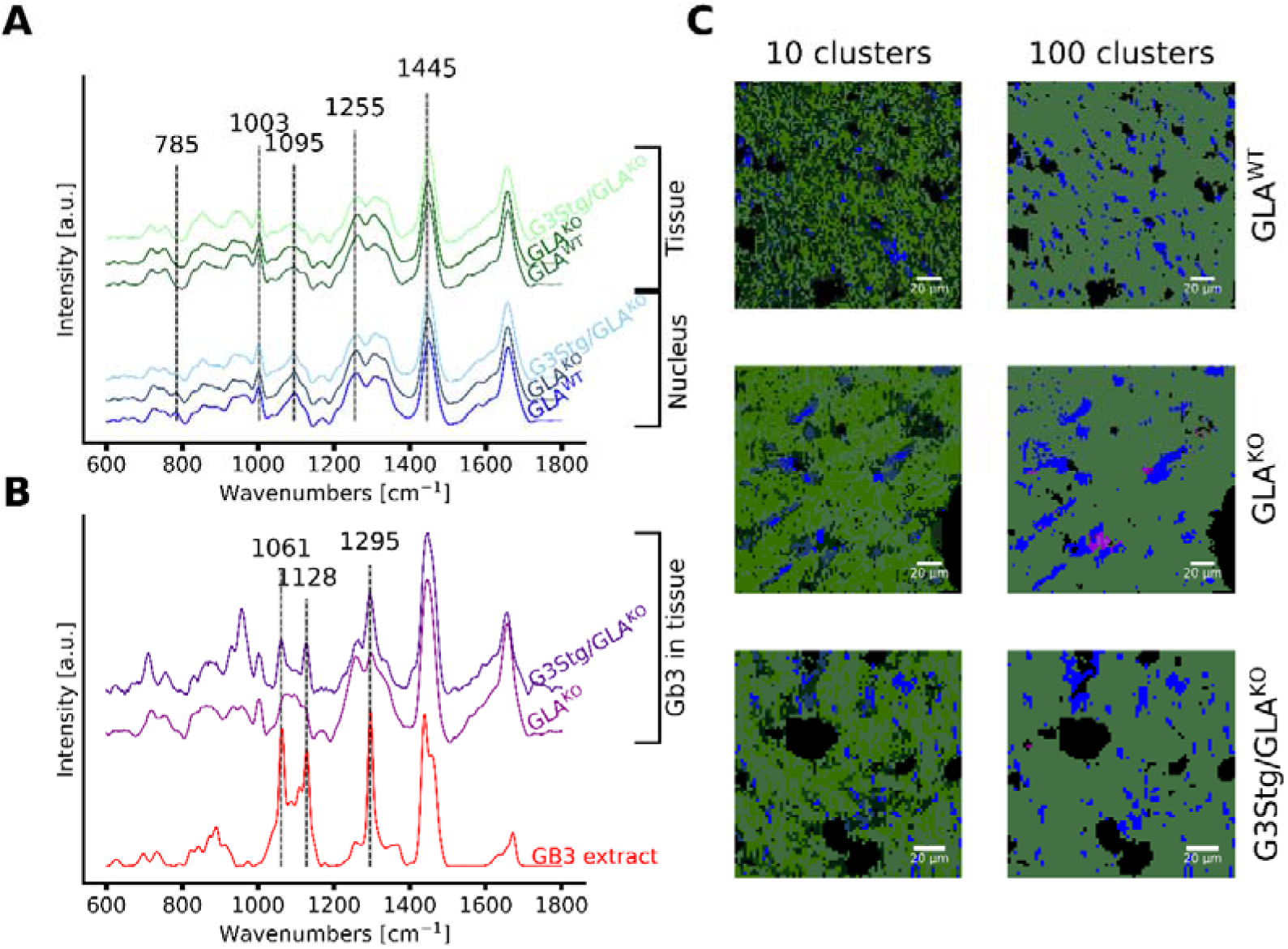
Multi-cluster hierarchical clustering analysis (HCA) of Raman imaging of tissue section. **A)** Mean spectra of the cluster analyses with 10 clusters. For each genotype, the mean spectrum of the nucleus and the mean spectrum of the remaining tissue were computed. Main intensity peaks (dashed lines) of nucleus and biological tissue components are visualized. Spectra are shifted along the y-axis for clarity. **B)** Mean spectra of clusters derived in HCA analyses with 100 clusters and associated with Gb3 compounds detected in both GLA^KO^ and G3Stg/GLA^KO^ samples (based on reference spectrum of commercial Gb3 mixture). **C)** Cluster images of corresponding HCA are shown for 10 and 100 clusters per genotype, respectively. Nuclei are depicted in blue; tissue is shaded in green,Gb3 is shaded in purple.

Aiming to distinguish additional chemical components within the tissue, the number of clusters was increased to 100. **Figure 4B** shows two additional clusters characterized by spectra with intense Raman peaks at 1061 cm^-1^, 1125 cm^-1^ and 1297 cm^-1^. Those peaks correlate with the characteristic intensity peaks of a commercial Gb3 extract of which a reference spectrum is shown in **Figure 4B**. The Gb3 clusters were localized only in a few pixels of tissue sections (**Figure 4C**, right column, in purple). Even though Raman peaks of the purified Gb3 extract and of certain pixels in the tissue sections correlate, identification of Gb3 by Raman imaging in tissue is challenging.

In order to characterize further molecular species using Raman imaging, we applied MCR-ALS to extract chemical components of Raman spectra and their corresponding relative abundance. The overview of all computed components found in the tissue is shown in the SI (**Figures S3-S4**). After MCR-ALS analysis, we selected four main components either based on their contribution to the overall signal or by biological relevance.

The distribution of the selected four components in the tissue of the three genotypes is shown in **Figure 5A** with corresponding spectra depicted in **Figure 5B**. The “nucleic acid” component revealed homogeneously distributed DNA spots in tissues with similar abundance throughout the three genotypes. The “tissue” component showed the morphological structure of the tissue, mainly based on the lipid/protein ratio. Additionally, the analysis identified a “collagen” component (intensity peaks at 858 cm^-1^, 947 cm^-1^ and 1240 cm^-1^)^61,62^, highlighting the ability of Raman imaging to detect biologically-relevant components that are not routinely captured with MALDI-MSI. Finally, the “lipid” component (intensity peaks at 1064 cm^-1^ and 1127 cm^-1^ (νC-C region), 1265 cm^-1^ (=CH), 1295 cm^-1^ (CH_2_) and 1439 cm^-1^ (CH_2_/CH_3_))^18,63,64^ revealed localized accumulations throughout the tissue with the highest abundance found in G3Stg/GLA^KO^. As the Gb3 clusters were not routinely separated from other components due to the spectral similarity with other lipids, a lipid-associated cluster was defined as a “lipids” group, shown in **Figure 5A**. Importantly, the lipid abundance measured by Raman was statistically significantly higher in G3Stg/GLA^KO^ cardiac tissue compared to the one of GLA^KO^ and GLA^WT^ mice, but not between GLA^KO^ and GLA^WT^.

**Figure 5.**
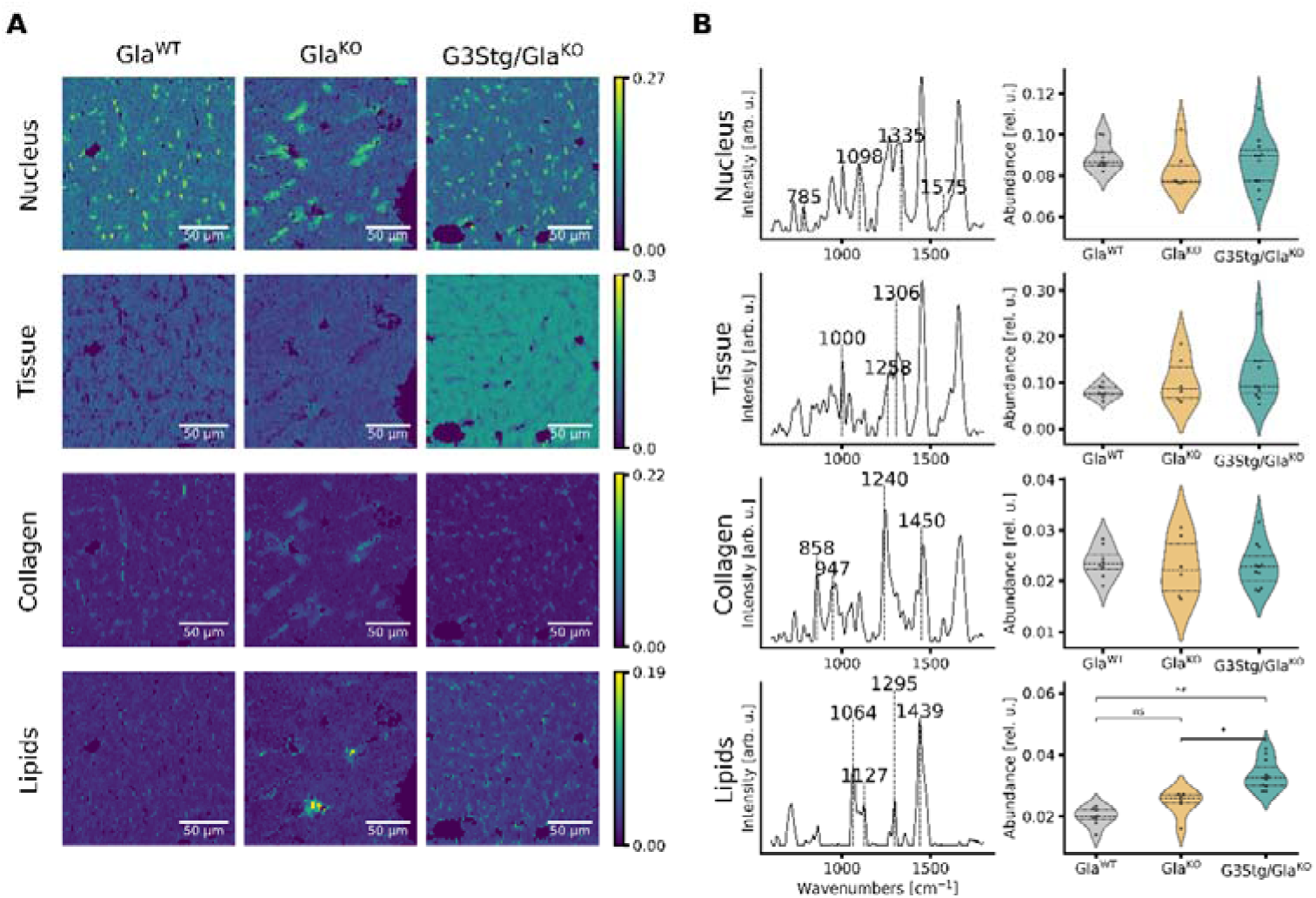
Component identification between different disease stages using MCR-ALS. **A)** The abundance maps of DNA-rich component (“nucleus”), a protein-rich component, (“tissue”), collagen, and lipids for a typical tissue example of GLA^WT^, GLA^KO^ and G3Stg/GLA^KO^. Images were acquired with 2 µm pixel resolution. Each intensity map was normalized to maximum intensity of this component over all measurements. **B)** The component spectra and the corresponding distribution of the mean abundance. The distribution plots show the mean summed intensity for each component. Statistical significance was tested on the mean intensity (derived from the mean abundance of the measurements) and visualized in case of significance. **, p < 0.01; *; p < 0.05; ns = not significant.

Raman imaging enabled a chemical component map of the cardiac tissue with various components: nucleic acids, lipid, protein, and collagen, and their relative abundance within the tissue. Whereas AP-MALDI MSI allowed the detection of the disease-specific lipid Gb3 and enabled the statistical differentiation between all three genotypes, Raman imaging offered the analysis and detection of a broader range of chemical components and revealed trends in overall lipid intensities that were consistent with MSI-based observations for Gb3 lipids.

### Automated co-registration of AP-MALDI-MS and Raman imaging of mouse heart tissue

Next, we aimed to integrate the complementary techniques, Raman imaging and AP-MALDI MSI at micrometre scale to maximize their combined analytical capabilities. Respective results are shown in **Figure 6**. For this integration, a software application for the automated co-registration of Raman imaging and AP-MALDI MSI datasets was developed. First, the Raman and MALDI-MSI results were overlaid with the corresponding BF images. For Raman imaging, BF was acquired at the same microscope enabling accurate identification of areas measured by Raman within the BF overview image. For MALDI MSI, BF images were acquired using an external microscope before and after each measurement. To map MALDI MSI frames onto the BF overview image acquired by the external microscope, we identified regions, where the MALDI laser had irradiated the tissue, as indicated by colour changes in the BF image. Thus, the areas on the tissue that displayed changes in the BF coloration were matched with the corresponding MALDI data. Subsequently, the BF-mapped Raman imaging and MALDI-MSI datasets were overlaid by spatially aligning the Raman- and MALDI-associated BF images. Examples of the individual steps involved in this process are shown in **Figure S5**.

**Figure 6.**
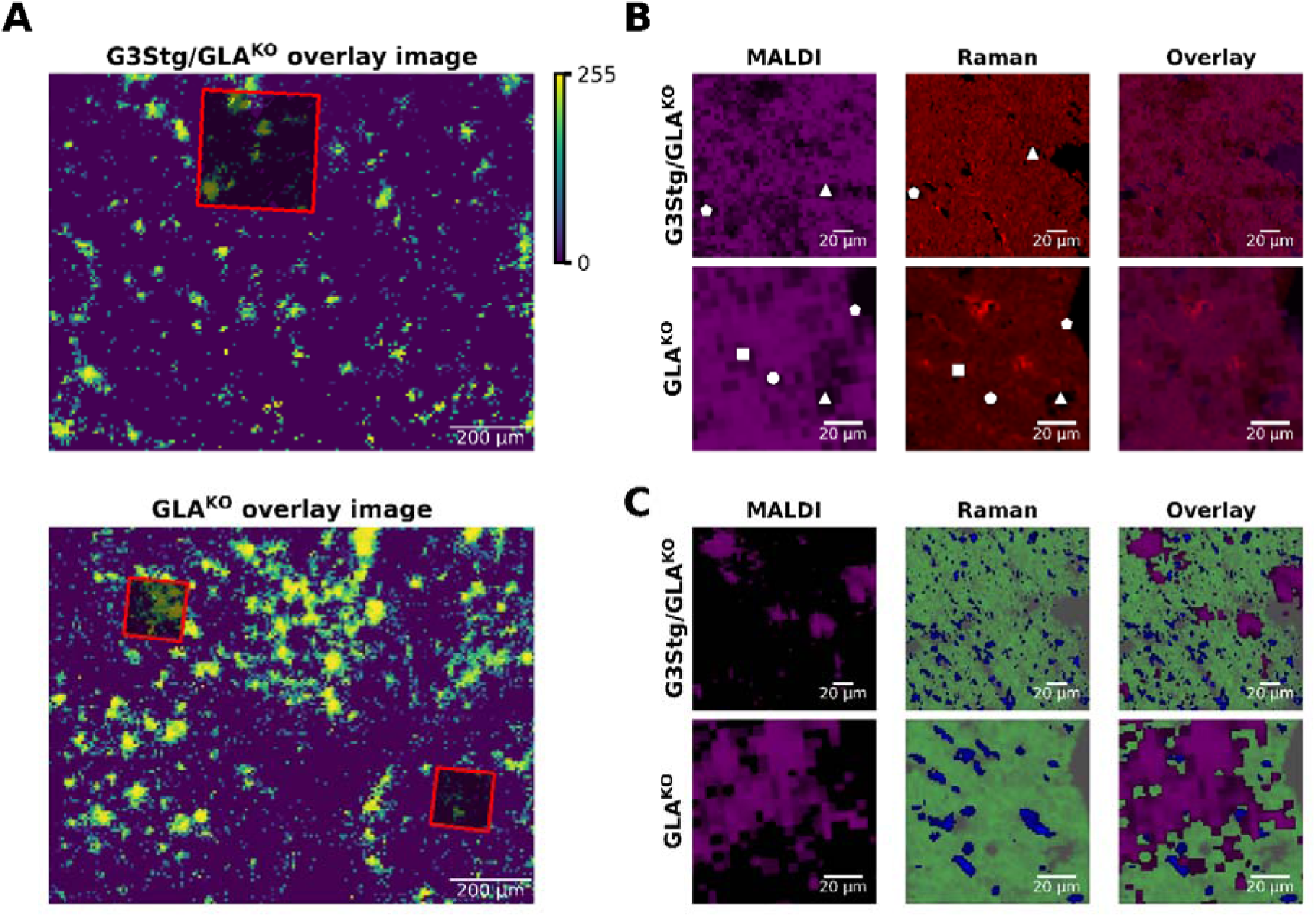
Automated overlay of AP-MALDI MSI and Raman images. **A)** RS measurements were located at their corresponding positions in the AP-MALDI MSI overview tissue measurement of Gb3(34:1)+Na]^+^ (*m*/*z* 1046.6591) for both G3Stg/GLA^KO^ and GLA^KO^ genotypes. **B)** [PC 34:1+Na]^+^ (*m*/*z* 782.5673) from AP-MALDI MSI results, Raman colour map of the lipid component and the overlay of both modalities. White symbols represent the same identified tissue features in both modalities. **C)** [Gb3(34:1)+Na]^+^ (*m*/*z* 1046.6591) accumulations (visualized in purple) from AP-MALDI MSI overlayed with “tissue” component in green and “nucleus” component in blue derived by MCR-ALS from Raman measurements. Scale: 20 µm.

In order to evaluate the alignment of Raman and MALDI MSI measurements, we focused on the three necessary tasks in the procedure: (1) the localization of Raman imaging results in the Raman-associated BF image (2) the localization of AP-MALDI MSI measurements in the MALDI-associated BF image, and (3) the co-localization of Raman- and MALDI-associated BF images. Tasks (1) and (2) were evaluated by calculating the Intersection over Union (IoU) between the computed rectangle coordinates and manually set reference boxes for both tasks.^65^ This resulted in IoU scores of 0.985 and 0.991 for AP-MALDI MSI and Raman imaging, respectively. These results indicate that automated localization of Raman imaging and MALDI MSI within corresponding BF images achieves comparable localization accuracy compared to manual area selection. For step (3), we evaluated the automatic co-localization performance by measuring distances between reference points in both BF images. This approach achieved an automated co-localization accuracy of (5.1 ± 1.6) µm for aligning Raman imaging and AP-MALDI MSI data, which is comparable to reported metrics for MALDI MSI localization methods in histopathological BF images^66^ but here enables the overlay of two complementary bioanalytical imaging techniques.

**Figure 6** illustrates co-registration results of Raman imaging and AP-MALDI MSI for both G3Stg/GLA^KO^ and GLA^KO^. In **Figure 6A**, Raman measurements (small red squares) are overlaid with the corresponding sodiated Gb3(34:1) overview images from AP-MALDI MSI.

To validate the co-localization performance of the automated overlay between Raman and AP-MALDI MSI, the distribution of the ubiquitous sodiated lipid PC 34:1 detected by AP-MALDI SI was compared to the lipid component distribution obtained from Raman imaging (**Figure 6B**). Due to partial tissue damage during sample preparation, certain regions lack tissue and consequently appear as areas with no signal intensity in both modalities. White symbols in **Figure 6B** highlight some of these positions. The presence and location of the same distinct features (white symbols in **Figure 6B**) in both Raman imaging and AP-MALDI MSI after automated co-localization confirms the validity of the approach within the reported error margins. Finally, we integrated the Gb3 distributions from AP-MALDI MS (visualized in pur**p**le) with additional biological components derived from Raman imaging, namely the cardiac tissue (green) and nucleic acids (blue) (**Figure 6C**). This multimodal data integration demonstrates the complementary analytical capabilities of both techniques. Raman imaging characterizes the morphological and biochemical structure of cardiac tissue by visualizing nuclei and tissue architecture including general lipids distribution, while AP-MALDI MSI specifically detects FD-related Gb3 accumulations enabling molecular differentiation.

### Conclusion and Outlook

We have developed a bioanalytical pipeline for the integration of AP-MALDI MSI and Raman spectroscopy at the low micrometre scale. While several seminal studies have combined Raman and MSI technologies,^23–26,67,68^ the results presented here represent the first integration of these two modalities that achieves co-localization of MSI data at 5 µm pixel size with Raman microscopy at 2 µm pixel size in pre-clinical mouse models.

We demonstrate that both methods independently capture significant changes in Gb3 and lipid accumulations, and that the combination of 5 µm - AP-MALDI MSI with Raman spectroscopy reveals the heterogonous landscape of Gb3 accumulations. In the GLA^KO^ and G3Stg/GLA^KO^ mouse FD models employed here, Gb3 accumulation within lysosomes and following lysosomal rupture in cardiac fibroblasts has been previously reported using fluorescence microscopy.^52,54^ The multimodal Raman/AP-MALDI MSI workflow developed in this study now provides increased molecular resolution for Gb3 accumulation in cardiac tissue. This approach revealed that Gb3 aggregation is not only heterogeneously distributed across the tissue but also exhibits variable lipidomic composition. This additional layer of heterogeneity among different Gb3 isoforms is not accessible by fluorescence microscopy methods. The automatic co-registration of AP-MALDI MSI, Raman imaging, and BF microscopy data with about 5 µm accuracy enabled contextualization of Gb3 accumulation within the tissue. Consequently, the presented AP-MALDI MSI method has the potential to significantly contribute to a better characterization of the heterogenous organization of Gb3 in tissues of the FD.

These developments will enable studies with larger cohorts that investigate the functional consequences of the molecular and spatial heterogeneity of Gb3 aggregation within cardiac tissue and thus rationalize the pathomechanisms underpinning these heterogeneities. Whereas AP-MALDI MSI revealed localized heterogenous regulation of Gb3 in heart tissue so far undetected via microscopic methods, Raman imaging provided tissue context but did not reveal new insights into the tissue architecture or molecular regulation in this setting. Therefore, future applications of this workflow will be expanded to human biopsy material to correlate different Gb3 lipoforms with patient phenotypes and evaluate whether therapeutic outcomes are associated with altered lipidomic profiles of Gb3 in cardiac tissue. Beyond FD, this pipeline has other potential applications in biomedical research where also Raman imaging can reveal biomolecular aggregation events, as for example the analysis of the microenvironment in myocarditis or diet-dependent accumulation of lipids in cardiac tissue.^69,70^

## Acknowledgement

All authors acknowledge the support by the “Ministerium für Kultur und Wissenschaft des Landes Nordrhein-Westfalen,” “Senatsverwaltung für Wissenschaft, Gesundheit und Pflege Berlin,” and by the “Bundesministerium für Forschung, Technologie und Raumfahrt” (BMFTR)”. The authors thank Felix-Levin Hormann, Chiahsin Chi, Ann-Katrin Weiss and Theresa Pietz for technical assistance and discussions.

## Supporting Information

H&E staining protocol, LC-MS/MS method description, LC-MS/MS results, AP-MALDI MSI results for Gb3 compound for different lipoforms, mean spectra and mean abundance of Raman results by MCR-ALS analysis for all genotypes, representative co-registration results.

## Notes

P. N. reveived advisory/speaker honoraria and research grants from Amicus, Chiesi, Idorsia, Sangamo, Sanofi, and Takeda. All other authors declare no competing financial interest.

## Author Contributions

Johann Dierks: Conceptualization, Investigation, Methodology, Data curation, Formal analysis, Writing, Review, Software, Validation, Visualization. Eike Brockmann: Conceptualization, Investigation, Methodology, Data curation, Formal analysis, Writing, Review, Validation, Visualization. Anahi-Paula Arias-Loza: Resources, Review. Thomas Bocklitz: Methodology, Investigation. Peter Nordbeck: Resources, Review. Kristina Lorenz: Conceptualization, Resources, Writing, Supervision, Project administration. Elena Tolstik: Conceptualization, Methodology, Investigation, Writing, Review, Supervision, Project administration. Sven Heiles: Conceptualization, Methodology, Investigation, Resources, Writing, Review, Supervision, Project administration.

