## Supporting Information for "Automatic colocalization of high resolution MALDI MSI and Raman imaging applied to cardiac tissue of Fabry disease mouse models"

¥ Shared first authors

§ Shared corresponding authors

### Materials and Methods

#### Post-acquisition H&E staining

Hematoxylin and eosin (H&E) staining of tissue sections following MALDI acquisition was performed according to a standardized protocol. Briefly, matrix-coated sections were washed in isopropanol for 2 minutes. Deparaffinization was then carried out using ROTI®Histol (6640.1, Carl Roth GmbH + Co. KG, Karlsruhe, Germany) in a three-step procedure, with each step lasting 4 minutes. Rehydration was performed sequentially using four washes of absolute ethanol and one wash of 70% ethanol, each for 2 minutes. Prior to staining, samples were rinsed in water four times, each rinse lasting 30 seconds.

Nuclear staining was conducted using Mayer’s hematoxylin (MHS32-1L, Sigma-Aldrich, Merck KGaA, Darmstadt, Germany) for 10 minutes, followed by a bluing step in running tap water for an additional 10 minutes. Counterstaining was performed with eosin G (yellowish) (1.15935.0100, Merck KGaA, Darmstadt, Germany; 10 g/L in Milli-Q water) for 50 seconds. Excess stain was removed by sequential rinsing in five water cuvettes.

Dehydration was achieved using 70% ethanol for 1 minute, followed by four successive washes in absolute ethanol, each lasting 2 minutes. Tissue clearing was performed with two changes of ROTI®Histol, 3 minutes each. Finally, stained sections were mounted using Eukitt® quick-hardening mounting medium (03989, Merck KGaA, Darmstadt, Germany) and allowed to dry overnight in a dark environment.

#### LC-MS/MS of cardiac tissue extracts for lipid identification

Lipid extracts were prepared from five to seven 20 µm-thick mouse heart sections per experimental group using a combined approach based on the Matyash (https://doi.org/10.1194/jlr.D700041-JLR200) and SIMPLEX (https://doi.org/10.1074/mcp.M115.053702) protocols. Samples were handled on ice whenever possible. All tissue sections from a given group were pooled into a single Eppendorf tube. Homogenization was carried out using the Bioruptor Pro (Diagenode(Details)) by adding 200 µL of ice cold 0.1% ammonium acetate (Details) in water(Details), 1 µL of SPLASH® Lipidomix® (Details), 1 µL of Gb3 C17:0 (100 µg/mL) (Details), and six to eight protein extraction beads (Details) to the tissue sample. The samples were sonicated at 4 °C using an alternating cycle of 30 seconds sonication and 30 seconds cooling, for a total duration of 10 minutes.

Lipid extraction was carried out by adding 120 µL of ice-cold methanol (MeOH (Details)) followed by 540 µL of ice-cold methyl tert-butyl ether (MTBE (Details)) to the homogenized samples. The mixture was incubated for 1 hour at 4 °C with continuous agitation at 1200 rpm. Phase separation was induced by the addition of 200 µL of 0.1% ammonium acetate in water. Samples were then centrifuged at 10,000 rpm for 10 minutes at 4 °C. The upper organic phase was carefully transferred to a clean Eppendorf tube. The remaining aqueous phase was subjected to a second extraction using the same procedure. Both organic phases were combined and the solvent was evaporated to dryness using a vacuum centrifuge (SpeedVac (Details)). The dried lipid extracts were reconstituted in 50 µL of methanol/isopropanol (1:1, v/v (Details)) and subsequently analyzed by liquid chromatography–tandem mass spectrometry (LC-MS/MS).

LC-MS/MS analysis was conducted using a Vanquish Flex UHPLC system (Thermo Fisher Scientific (Details)) coupled to an Orbitrap Exploris 240 mass spectrometer (Thermo Fisher Scientific (Details)). Chromatographic separation was achieved on an Ascentis® Express C18 column (150 × 2.1 mm, 2.7 µm particle size, 90 Å pore size; Supelco, Germany) using a 36-minute stepped gradient from 10% to 100% mobile phase B.

Mobile phase A consisted of acetonitrile/water (1:1, v/v (Details)) supplemented with 5 mM ammonium formate (Details) and 0.1% formic acid (Details). Mobile phase B consisted of isopropanol/acetonitrile/water (85:10:5, v/v/v) with 5 mM ammonium formate and 0.1% formic acid. The gradient profile was as follows: 0–20 min, 10–86% B; 20–22 min, 86–100% B; 22–27 min, 100% B; 27–28 min, 100–10% B; 28–36 min, 10% B. The flow rate was set to 300 µL/min, and the column temperature was maintained at 50 °C. The injection volume was set to 5 µL.

Mass Spectrometric Parameters were as follows: The total method duration was 34 minutes. The H-ESI parameters were configured as follows: a static spray voltage of 3.1 kV, sheath gas flow rate of 40 (arbitrary units), auxiliary gas flow rate of 15, and sweep gas flow rate of 12. The ion transfer tube was maintained at a temperature of 320 °C, while the vaporizer temperature was set to 300 °C. The expected LC peak width was set to 30 seconds, and advanced peak determination was enabled. The default charge state was assigned as +1, and lock mass correction was applied using the EASY-ICTM system.

Full-scan mass resolution was set to 180,000, and for data-dependent MS/MS (ddMS2) scans, a resolution of 15,000 was employed, covering an m/z range of 150–1200. The RF lens level was adjusted to 70%. The automatic gain control (AGC) target was set to standard, and the maximum injection time was operated in automatic mode. Data acquisition was performed in positive ionization mode with profile detection, utilizing one microscan and no source-induced fragmentation. The intensity threshold for precursor selection was set to 1000.

Dynamic exclusion was enabled in custom mode, with exclusion applied after a single occurrence and maintained for 15 seconds. The mass tolerance for precursor selection was ±5 ppm for both high and low masses, and isotopic peaks were excluded. Data-dependent acquisition (DDA) was configured to collect ten MS/MS scans per cycle, employing an isolation window of m/z 1.5 and normalized higher-energy collisional dissociation (HCD) collision energies of 20%, 25%, and 30%.

### Results

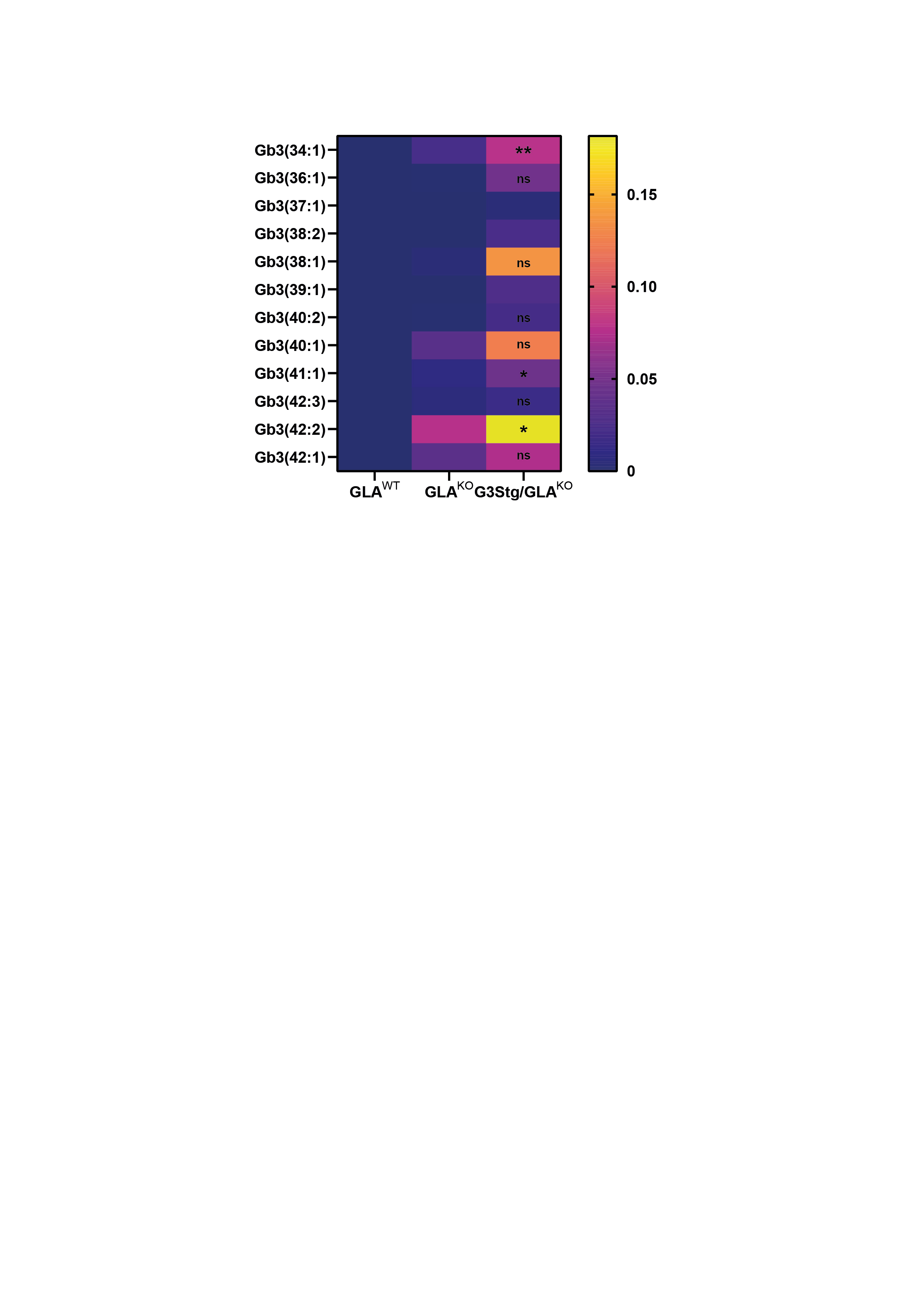

#### **Figure S1:** Heatmap of normalized mean ROI intensities:

The heatmap demonstrates the mean intensity differences within the groups for all Gb3 species as detected with AP-MALDI-MSI. The displayed intensities represent mean ROI intensities of the tissue area and were calculated following hot-spot removal by histogram equivalation and RMS normalization. GLA^WT^ n=5, GLA^KO^ n=4, G3Stg/GLA^KO^ n=3 biological replicates. ** p < 0.01; * p < 0.05; ns = not significant. Significances were calculated as described in the methods section.

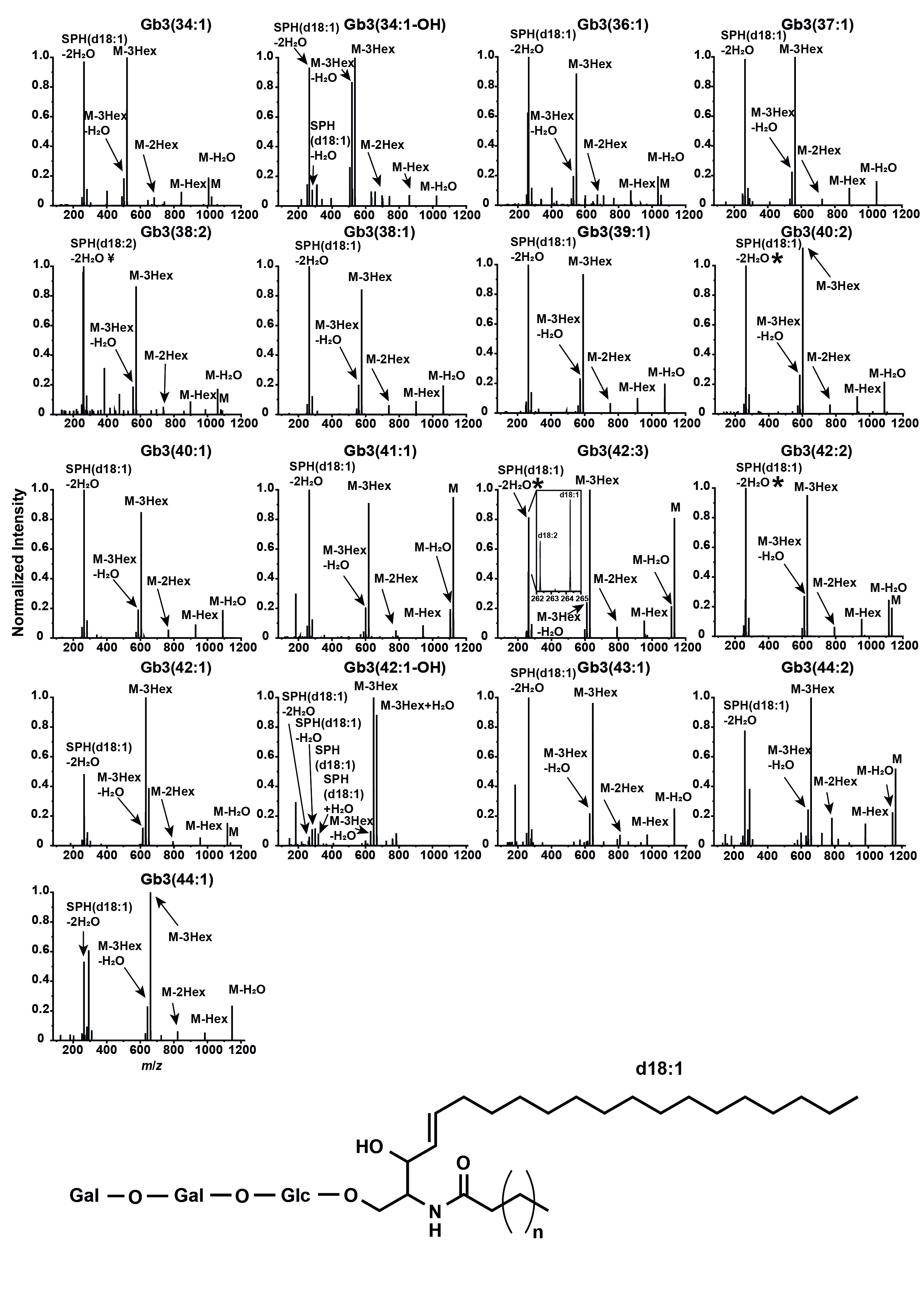

#### **Figure S2:** LC-MS/MS fragment spectra of all detected Gb3 species.

HCD of protonated Gb3 species resulted in distinct fragment spectra with a sequential loss of the three hexoses (Hex) and the additional cleavage of the fatty acyl chain and double water loss, resulting in the sphingosine moiety at m/z 264.26. * For Gb3(40:2), Gb3(42:2) and Gb3(42:3) the sphingadiene moiety (*m*/*z* 262.26) could be detected in addition to m/z 264.26. For Gb3(38:2), only the sphingadiene moiety could be detected.

**Table S1:** List of detected Gb3Cer species as identified by accurate mass in AP-MALDI MSI and LC-MS/MS.

| **Gb3-Species** | **Sphingosine** | **Fatty acid** | **Exact mass [M+H]^+^** | **Δ (mDa)** | **ppm** | **LC-MS/MS** | **AP-MALDI-MSI [M+Na]^+^, [M+K]^+^** |
| --- | --- | --- | --- | --- | --- | --- | --- |
| **Gb3(d18:1/C16:0)** | d18:1 | C16:0 | 1024.6778 | 0.58 | 0.6 | x | x |
| **Gb3(d18:1/C16:0+OH)** | d18:1 | C16:0-OH | 1040.6727 | 0.58 | 0.6 | x | n/a |
| **Gb3(d18:1/C18:0)** | d18:1 | C18:0 | 1052.7091 | 0.18 | 0.2 | x | x |
| **Gb3(d18:1/C19:0)** | d18:1 | C19:0 | 1066.7248 | 0.32 | 0.3 | x | x |
| **Gb3(d18:2/C20:0)** | d18:2 | C20:0 | 1078.7248 | 0.58 | 0.5 | x | x |
| **Gb3(d18:1/C20:0)** | d18:1 | C20:0 | 1080.7404 | 1.02 | 0.9 | x | x |
| **Gb3(d18:1/C21:0)** | d18:1 | C21:0 | 1094.7561 | 1.02 | 0.9 | x | x |
| **Gb3(d18:2/C22:0)** | d18:2 | C22:0 | 1106.7561 | 0.22 | 0.2 | x | n/a |
| **Gb3(d18:1/C22:0)** | d18:1 | C22:0 | 1108.7717 | 0.52 | 0.5 | x | x |
| **Gb3(d18:1/C22:1)** | d18:1 | C22:1 | 1106.7561 | 1.22 | 1.1 | x | x |
| **Gb3(d18:1/C23:0)** | d18:1 | C23:0 | 1122.7874 | 0.32 | 0.3 | x | x |
| **Gb3(d18:2/C24:0)** | d18:2 | C24:0 | 1134.7874 | 0.82 | 0.7 | x | n/a |
| **Gb3(d18:1/C24:0)** | d18:1 | C24:0 | 1136.8030 | 0.12 | 0.1 | x | x |
| **Gb3(d18:1/C24:0+OH)** | d18:1 | C24:0-OH | 1152.7979 | 0.28 | 0.2 | x | - |
| **Gb3(d18:2/C24:1)** | d18:2 | C24:1 | 1132.7718 | 1.02 | 0.9 | x | n/a |
| **Gb3(d18:1/C24:1)** | d18:1 | C24:1 | 1134.7874 | 0.82 | 0.7 | x | x |
| **Gb3(d18:1/C24:2)** | d18:1 | C24:2 | 1132.7717 | 1.12 | 1 | x | x |
| **Gb3(d18:1/C25:0)** | d18:1 | C25:0 | 1150.8187 | 0.12 | 0.1 | x | - |
| **Gb3(d18:1/C26:0)** | d18:1 | C26:0 | 1164.8343 | 0.78 | 0.7 | x | - |
| **Gb3(d18:1/C26:1)** | d18:1 | C26:1 | 1162.8187 | 0.30 | 0.2 | x | - |

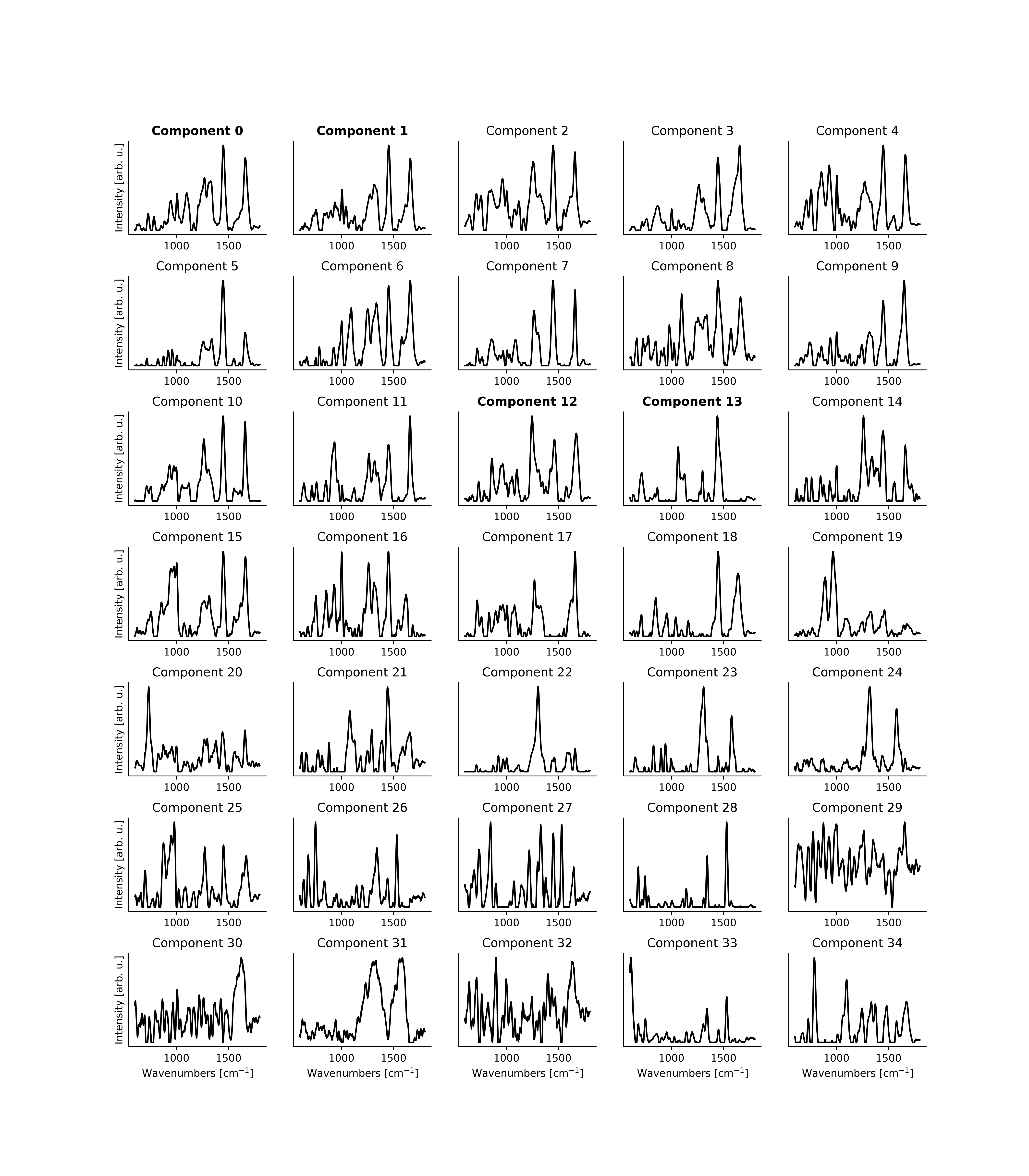

#### **Figure S3:** Mean spectra for all computed 35 components found in the tissues and identified using MCR-ALS.

The components are sorted by frequency of occurrence in the tissue. From all 35 components only four main components were selected for further study either based on their contribution to the overall signal or by biological relevance, see also **Figure S4**.

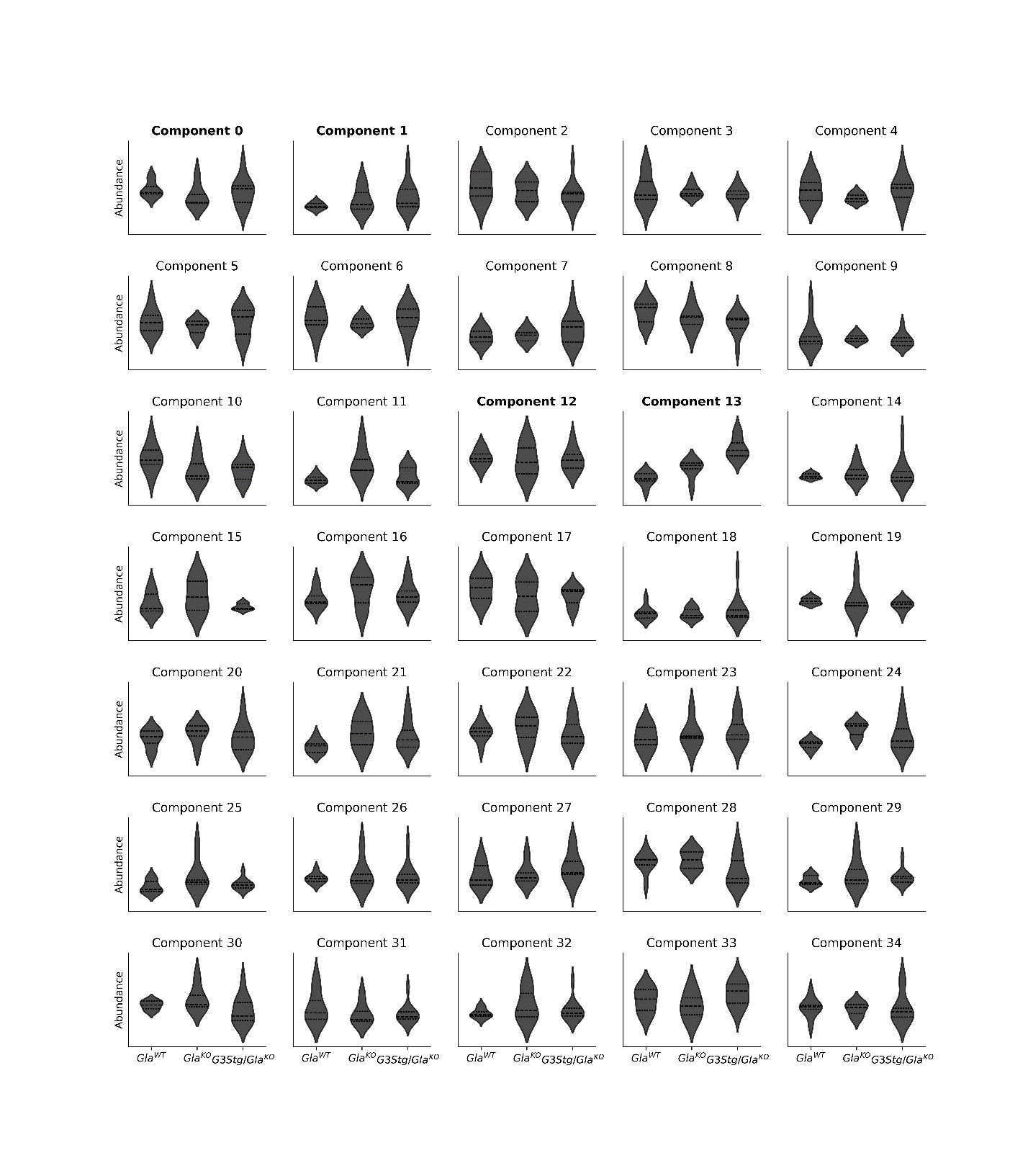

#### **Figure S4:** Distribution of the mean abundance for all computed components found in the tissues and identified using MCR-ALS.

The components are sorted by frequency of occurrence and each component is presented for three genotypes: GLA^WT^, GLA^KO^ and G3Stg/GLA^KO^. From all 35 components only four main components were selected for further study either based on their contribution to the overall signal or by biological relevance.

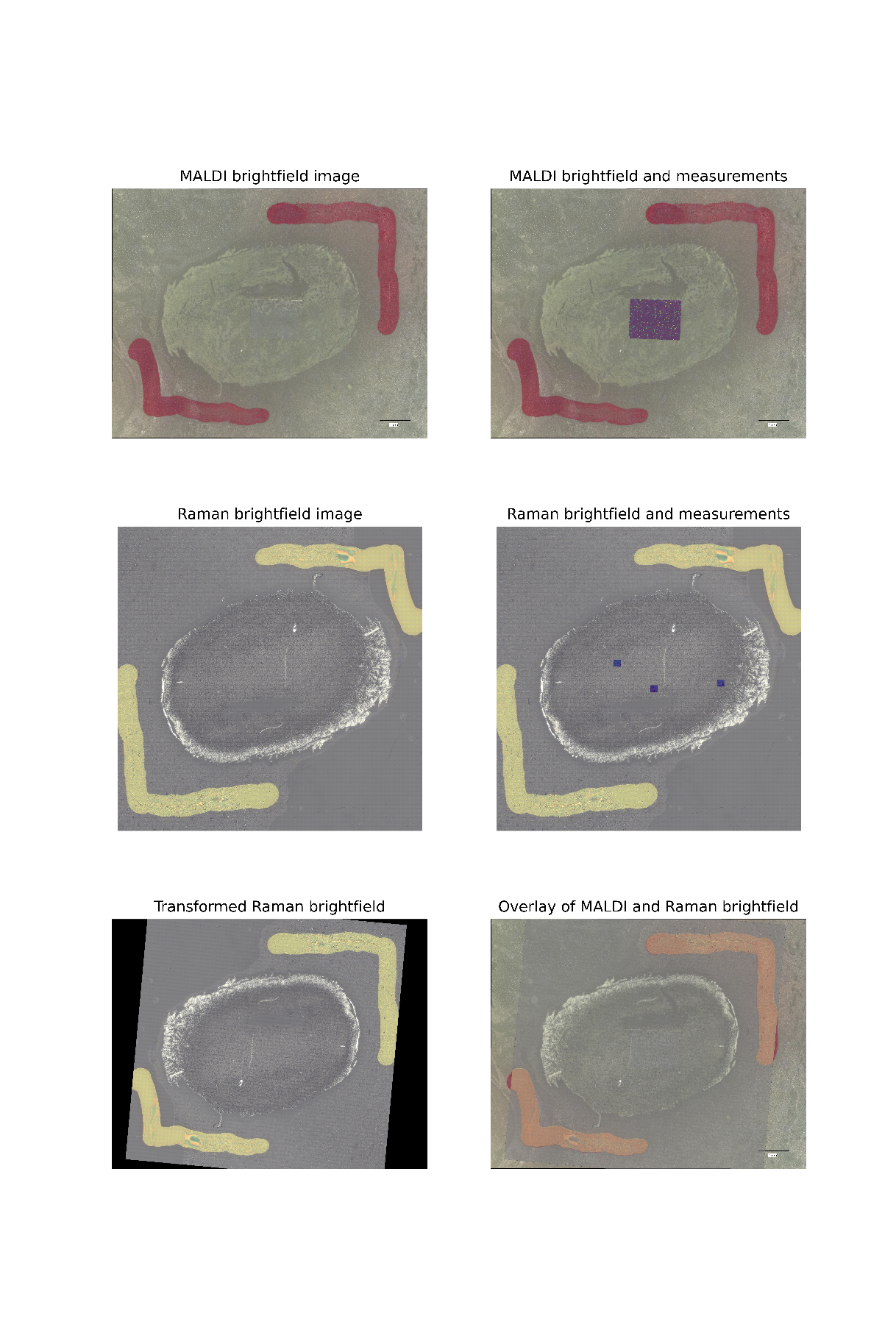

#### **Figure S5:** Example of the superposition between MALDI, Raman and BF imaging.

The two overview BF images are labelled “MALDI brightfield image” and “Raman brightfield image”. Localization of MALDI and Raman results in the BF images, as described for defined task (1) and (2), are shown on the right and are labelled “MALDI brightfield and measurement” and “Raman brightfield and measurement”, respectively. Coalignment of both BF images, task (3), and therefore coalignment of MALDI and Raman results, are shown in the lower row and are labelled “Transformed Raman brightfield” and “Overlay of MALDI and Raman brightfield”.
